# An AI-driven pipeline for the discovery of hidden peptides in plant proteomes: the CLE family as a case study

**DOI:** 10.64898/2026.01.31.703007

**Authors:** Marco Boschin, Marco Rota Negroni, Carlotta Francese, Anna Pavanello, Gabriele Sales, Livio Trainotti

## Abstract

Plant proteomes contain evolutionarily conserved peptides with poorly conserved primary sequences, often hindering their identification and classification into families. Homology-based approaches and conventional annotation pipelines frequently fail to detect these family members, particularly in poorly characterized, but agronomically relevant plant species. CLE peptides (CLAVATA3/EMBRYO SURROUNDING REGION–related peptides) constitute a large and evolutionarily conserved family of plant signaling molecules, yet their characterization remains incomplete. Beyond a limited number of well-studied members, a substantial number of CLE peptides remain uncharacterized due to functional redundancy and the intrinsic features of *CLE* genes, which encode short pre-propeptides with only a small 12-residue conserved motif.

Here, we present a novel framework leveraging state-of-the-art Protein Language Models (pLMs) to discover CLE peptides directly from 13 plant proteomes. By coupling sequence embeddings trained on large evolutionary datasets (ESM2 and ProtT5) with supervised machine learning, our dual-model approach captures deep semantic features of the CLE family that are missed by traditional alignment methods. The pipeline demonstrated robust generalization, achieving high classification accuracy (98.9-99.4%) on a held-out set of CLE peptides not used during training. Consequently, we identified a set of high-confidence, previously unannotated CLE candidates prioritized through a stringent consensus-based filtering strategy.

This work demonstrates how AI-driven proteome analysis can overcome the limitations of homology-based methods and provides a scalable strategy for uncovering previously unidentified peptide-mediated signaling molecules across plant lineages.

**Highlight:** Leveraging Protein Language Models, our AI framework uncovers “hidden” signaling peptides missed by standard tools, revealing the elusive diversity of CLE regulators across plant proteomes.

## Introduction

CLE peptides (CLAVATA3/EMBRYO SURROUNDING REGION-related peptides) are small signaling peptides of approximately twelve amino acids, initially identified for their role in controlling meristematic activity.

The discovery of CLAVATA3 (CLV3) in *Arabidopsis thaliana* represented a turning point in the study of peptide-mediated communication in plants, demonstrating how endogenous peptide signals finely regulate the maintenance of the shoot apical meristem and the balance between stem cells and differentiated cells (Clark *et al*., 1995; Fletcher *et al*., 1999). After the identification of *CLV3*, the search for orthologous genes in other species led to the discovery of the EMBRYO SURROUNDING REGION (ESR) proteins in *Zea mays* (Opsahi-ferstad *et al*., 1997) and, subsequently, of numerous other peptides sharing the same bioactive sequence as CLV3. These peptides were collectively designated as CLE (CLAVATA3/ESR-related). Numerous members of this family have been identified across a wide range of plant species, revealing an extraordinary diversity of biological functions.

Besides meristem development, CLE peptides are now known to regulate processes such as phloem and xylem formation, nodulation, responses to abiotic stress, leaf senescence and fruit development, among others (Yamaguchi *et al*., 2016; Fletcher, 2020; Cornelis and Hazak, 2025).

*CLE* genes are relatively small and encode non-functional pre-propeptides of approximately 100 amino acids, which contain an N-terminal signal peptide, a central variable region, and a highly conserved C-terminal CLE domain (Yamaguchi *et al*., 2016; Fletcher, 2020). The pre-propeptide undergoes proteolytic processing by serine-proteases and carboxypeptidases at positions flanking the CLE region, resulting in the release of the mature, biologically active CLE peptide (Djordjevic *et al*., 2011; Ni *et al*., 2011; Goad *et al*., 2017). In their route through the secretory pathway, CLE peptides also go through post-translational modifications; in particular, proline hydroxylation (at P4 and/or P7) and arabinosylation (at Hyp-7) strengthen the interaction between the peptide and its receptor (Yamaguchi *et al*., 2016; Fletcher, 2020; Stührwohldt *et al*., 2020).

Once processed, the CLE peptides are secreted into the apoplast, where they function as signaling molecules by binding to leucine-rich repeat receptor-like kinases (LRR-RLKs), such as CLV1, or receptor-like proteins (RLPs), like CLV2, thereby facilitating the transmission of extracellular signals across the membrane (Hohmann *et al*., 2017) and triggering downstream signaling cascades (Olsson *et al*., 2019).

The specificity of peptide–receptor interactions can vary depending on the peptide involved. A single receptor may bind multiple peptides, and conversely, an individual peptide can be recognized by more than one receptor (De Coninck and De Smet, 2016). In most cases, high-affinity binding between peptide ligands and their receptors requires the presence of a co-receptor (Hu *et al*., 2018). Ligand binding typically induces conformational changes in the receptor (Zhang *et al*., 2016), which can trigger the recruitment of co-receptors and initiate rapid biochemical events, including receptor (auto)phosphorylation, protein–protein interactions, and phosphorylation of downstream targets. These early molecular events activate signaling cascades that ultimately lead to specific cellular responses.

The expansion of the *CLE* gene family across plant genomes has resulted in a remarkable diversity of peptide members with different functions. This functional diversity arises through different evolutionary mechanisms, including whole-genome and segmental duplication. Moreover, the specificity of *CLE* functions is primarily achieved through differential expression patterns (subfunctionalization at the level of regulatory sequences) rather than through biochemical divergence (neofunctionalization) (Jun *et al*., 2007; Gentile *et al*., 2025). Based on current literature, CLE peptides are widely distributed across all major groups of land plants, whereas no CLE homologs have been identified in algae, indicating that this peptide family likely originated in the last common ancestor of land plants (Whitewoods *et al*., 2018; Whitewoods, 2021). Mosses present a notable exception regarding the TDIF-like subclass of CLE peptides. In vascular plants, TDIF-like CLEs play a key role in vascular development, specifically by inhibiting the differentiation of tracheary elements. However, contrary to other bryophytes, mosses, which naturally lack vascular tissues, have secondarily lost this subclass during evolution, likely reflecting the absence of the corresponding developmental context (Whitewoods, 2021).

Since the discovery of *CLV3*, extensive efforts have been devoted to identifying paralogous genes. In the earliest investigations, this challenge proved difficult due to the short length and high sequence variability of these peptides (Cock and McCormick, 2001). Nevertheless, the identification of CLE peptides has evolved substantially over time. Early studies relied primarily on sequence homology searches. These initial analyses led to the identification of 42 sequences from seven different plant species; in *A. thaliana* 24 genomic sequences were identified (Cock and McCormick, 2001). Subsequent analyses, together with experimental validation, have updated this number, and CLE peptides are currently considered to comprise 32 members in *A. thaliana*, most of which were reported to be expressed across one or more organ in different tissues (Hobe *et al*., 2003; Sharma *et al*., 2003; Fiers *et al*., 2006; Strabala *et al*., 2006; Wang and Fiers, 2010). Over the years, increasingly sensitive probabilistic methods have been introduced. In particular, Hidden Markov model (HMM)–based strategies have enabled a more comprehensive detection of *CLE* genes, as exemplified by iterative BLAST and HMM-based pipelines that substantially expanded the CLE repertoire in crop species. An illustrative example is represented by the discovery of CLE peptides in the tomato genome. Until 2014, only 15 CLE peptides had been reported in *Solanum lycopersicum* (Zhang *et al*., 2014). Later, Carbonnel *et al*., (2022) significantly expanded the dataset through genome-wide analysis, bringing to 52 the total number of *CLE* genes in tomato. Nowadays, the field of peptide discovery has entered a new phase driven by large-scale genomics. Indeed, interesting research studies have been carried out at pan-angiosperm level, combining *de novo* genome annotation, machine learning–based sequence embeddings, and protein language models, which have enabled the discovery of thousands of previously unannotated CLE peptides across deep evolutionary timescales, thereby overcoming the intrinsic limitations of homology-dependent methods (Gentile *et al*., 2025).

As CLE peptides were uncovered in an increasing number of plant genomes, it became clear that some *core* CLE peptides function as key regulators of fundamental developmental processes. The functional predominance of these key players can sometimes obscure the potential roles of many other family members, which are likely involved in the fine-tuning of numerous physiological processes.

In addition, the very short CLE peptide length and low sequence conservation outside the CLE domain make *CLE* genes particularly difficult to identify using general genome annotation tools. This leads to underrepresentation of CLEs in functional studies, especially in poorly characterized species that nonetheless play an important role in agricultural production. As a consequence, the biological relevance of many CLE peptides remains largely unexplored, representing a significant gap in our understanding of CLE-mediated signaling in plants.

CLE peptides do not act in isolation, but function within interconnected signaling networks, in which multiple families of small secreted peptides collectively regulate plant development and physiology. Investigating CLE peptides outside their biological context provides only a partial view of their function, as developmental outcomes arise from the integration of diverse signaling inputs. Indeed, the ROOT GROWTH FACTOR/CLE-LIKE/GOLVEN (RGF/CLEL/GLV) family was identified after the CLE peptides (Matsuzaki *et al*., 2010), but shares key structural features with CLEs, including overlapped biogenesis and post-translational processing leading to the secretion of active signals (Olsson *et al*., 2019). GLV peptides display distinct molecular properties, such as differences in peptide length and conserved sequence features, which distinguish them from canonical CLE peptides and are consistent with specialized biological functions. GLVs are well known for their role in root development, lateral root formation, and root meristem regulation (Meng *et al*., 2012).

Together, these observations underscore the importance of adopting a network-oriented perspective when studying CLE peptides, in which their functions are interpreted within the broader landscape of peptide-mediated signaling rather than as isolated regulatory elements. Thus far, most studies have focused on genomic analyses (Zhang *et al*., 2020; Carbonnel *et al*., 2022), while less attention has been paid to proteomic analyses, which reflect proteins that are actually expressed in tissues. To overcome the limitations of homology-dependent methods in characterizing these elusive signaling molecules, we propose a novel artificial intelligence–based approach for the identification of regulatory peptides in previously unexplored proteomes.

In recent years, protein language models (pLMs) have emerged as transformative tools in computational biology. These models learn rich numerical representations of protein sequences, termed embeddings, by training on hundreds of millions of unlabeled sequences. The fundamental insight is that protein sequences, like natural language, exhibit contextual dependencies: the identity of each amino acid is constrained by its surrounding sequence context. By learning to predict masked amino acids from their neighbors, pLMs internalize the underlying “grammar” of proteins, capturing evolutionary and structural relationships without explicit annotation (Rives *et al*., 2021).

The resulting embeddings encode proteins as points in a high-dimensional vector space. As described by Hayes *et al*. (2025), “proteins can be seen as existing within an organized space where each protein is neighbored by every other protein that is one mutational event away.

The structure of evolution appears as a network within this space, connecting all proteins by the paths that evolution can take between them.” In this learned representation, functionally related proteins cluster together regardless of overall sequence identity, while functionally divergent proteins occupy distant regions, enabling the detection of remote homologs that escape traditional sequence alignment methods. Because a language model is trained to predict the next token, it must learn the landscape of evolutionary possibilities within the space of possible proteins, effectively encoding millions of years of evolutionary constraints into its representations.

This property makes pLMs particularly suited for identifying small signaling peptides like CLEs, where the conserved functional domain comprises only ∼12 amino acids within larger, variable precursors (70–150 residues). Unlike traditional homology-based approaches, which struggle with such short, rapidly evolving sequences, pLM embeddings capture both the conserved core and contextual information, enabling identification of CLE precursors even without detectable sequence similarity to known family members.

Building on these principles, for this study we employed a strategy combining two architecturally distinct pLMs: ESM2 (Lin *et al*., 2023), a leading BERT-like encoder model, and ProtT5 (Elnaggar *et al*., 2022), based on the encoder-decoder T5 architecture. These models employ different embedding strategies (Yeung *et al*., 2023), and were trained with distinct objectives, capturing complementary aspects of protein sequence information. By combining large-scale proteomic screening with this AI-driven framework (***Fig. 1A***), we introduce a robust pipeline for uncovering novel CLE peptides, with a methodology that is potentially generalizable to other protein families.

**Fig. 1.**
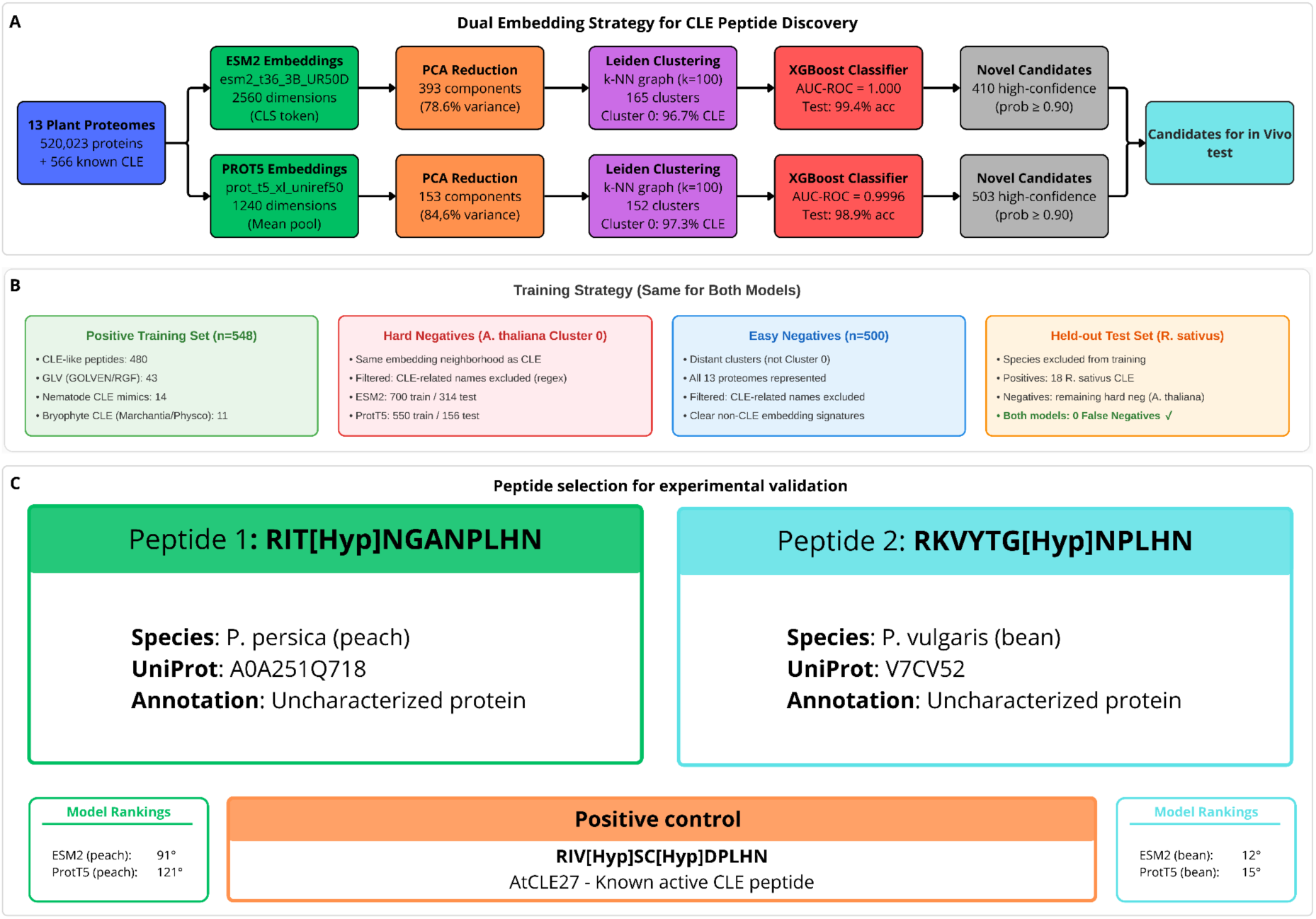
AI-driven pipeline for CLE peptide discovery and experimental validation strategy. (**A**) Overview of the dual-embedding workflow. The pipeline processes 520,023 protein sequences from 13 plant proteomes using two independent Protein Language Models (ESM2 and ProtT5). High-dimensional embeddings are reduced via PCA and clustered using the Leiden algorithm. XGBoost classifiers are trained on the CLE-enriched clusters to identify high-confidence candidates (probability ≥0.90). (**B**) Training set composition. The strategy employs a stratified approach including known CLEs (positives), hard negatives (non-CLEs from the CLE-enriched cluster), and easy negatives (distant proteins). A held-out test set of *R. sativus* CLEs was used to validate model generalization. (**C**) Candidate selection for *in vivo* testing. Two high-confidence consensus candidates were selected: PEP1 (*P. persica*) and PEP2 (*P. vulgaris*). Both peptides rank highly in both models and share partial sequence similarity with known CLEs while lacking prior functional annotation. *At*CLE27 is included as a positive control.

## Materials and methods

### Dataset assembly

Complete proteomes were obtained from UniProt (The UniProt Consortium, 2023) for 11 plant species: *Arabidopsis thaliana* (UP000006548), *Solanum lycopersicum* (UP000004994), *Prunus persica* (UP000006882), *Vitis vinifera* (UP000009183), *Zea mays* (UP000007305), *Glycine max* (UP000008827), *Cucumis melo* (UP000089565), *Phaseolus vulgaris* (UP000000226), *Marchantia polymorpha* (UP000244005), *Physcomitrium patens* (UP000006727), and *Raphanus sativus* (UP000504610). *Fragaria vesca* (v4.0.a1) and *Lotus japonicus* (Lj1.0v1) proteomes were obtained from GDR (Jung *et al*., 2019) and Phytozome (Goodstein *et al*., 2012), respectively. *Marchantia polymorpha* and *Physcomitrium patens* were included as phylogenetically distant species (bryophytes) to assess whether the embedding space captures conserved CLE features across deep evolutionary divergence. The combined dataset comprised 520,023 protein sequences.

Known CLE and CLE-related peptides were compiled from UniProt annotations, published literature (Hastwell *et al*., 2015, 2017; Gancheva *et al*., 2016; Goad *et al*., 2017; Whitewoods *et al*., 2018; Hirakawa *et al*., 2019; Wang *et al*., 2019; Qin *et al*., 2021; Carbonnel *et al*., 2022) and unpublished data previously generated in our laboratory. The positive training set included 548 sequences: 480 CLE-like peptides from multiple species, 43 GLV (RGF/CLEL/GLV) peptides ((Fang *et al*., 2021), and laboratory unpublished data), 14 nematode-derived CLE mimics obtained from *Heterodera glycines*, *H. schachtii*, *Globodera rostochiensis*, and *Rotylenchulus reniformis* (Lu *et al*., 2009; Wang *et al*., 2011; Wubben *et al*., 2015) and 11 CLE peptides from *Marchantia polymorpha* and *Physcomitrium patens*. *Raphanus sativus* CLE sequences (n=18) were reserved as a test set to evaluate model generalization.

### Generation of protein embeddings

To ensure robust candidate identification, we employed two independent pLMs with distinct architectures and training objectives. We first ran ESM2 (Lin *et al*., 2023), specifically the esm2_t36_3B_UR50D model with 3 billion parameters. We extracted the CLS token representation for each sequence from the final transformer layer, yielding 2560-dimensional embeddings.

We then used the ProtT5-XL-UniRef50 model (Elnaggar *et al*., 2022), a protein-specific encoder-decoder based on the T5 architecture. Sequence representations were obtained by mean pooling over all residue-level hidden states from the final encoder layer, producing 1024-dimensional embeddings.

Both embedding sets were processed in parallel through identical downstream pipelines. All computations were performed on a NVIDIA H100 GPU.

### Dimensionality reduction and clustering

For each embedding type, Principal Component Analysis (PCA) was applied to reduce dimensionality. The optimal number of components was determined independently using the Kneedle algorithm (Satopaa *et al*., 2011) on the cumulative explained variance curve. For ESM2 embeddings, this yielded 393 principal components (78.6% variance explained); for ProtT5, 153 components (84.6% variance explained).

Clustering stability was assessed following (Zappia and Oshlack, 2018). A subsample of 50,000 sequences was used to test k-NN graph construction with k values of 10, 15, 20, 30, 50, and 100. For each value of *k*, 20 iterations of 80% subsampling were performed, and clustering consistency was measured using the Adjusted Rand Index (ARI) between independent subsamples. Both embedding types showed optimal stability at k=100, with median ARI of 0.755 (ESM2) and 0.729 (ProtT5).

Community detection was performed using the Leiden algorithm (Traag *et al*., 2019) on k-NN graphs with modularity optimization. Clustering was performed on PCA-reduced coordinates. UMAP and t-SNE transformations were used for visualization purposes.

### Cluster enrichment analysis

We computed the fraction of known CLE-related peptides within each cluster. Clusters were considered enriched if they met the following criteria: ≥5% positive peptides, an absolute count of ≥10 positive sequences, and entropy ≤1.5. Entropy was calculated as Shannon entropy based on the relative frequency of positive sequence categories within each cluster; this threshold was applied to ensure that positives were concentrated within coherent functional groups rather than dispersed across many, reflecting a robust biological signal. We confirmed that clustering reflected functional rather than taxonomic relationships by ensuring no cluster was dominated by a single species (containing >80% sequences from a single proteome).

### XGBoost classification

Independent XGBoost classifiers were trained on each embedding type to discriminate CLE peptides from non-CLE proteins. We employed a stratified negative sampling strategy divided into two categories to maximize model robustness (***Fig. 1B***). We considered *A. thaliana* proteins located within the CLE-enriched cluster (Cluster 0) which were lacking CLE-related annotation as ‘hard negative’ examples. We focused on this species due to its comprehensive proteome annotation, which reduced the risk of ‘label noise’ (unannotated true CLEs in the negative set). We focused on this species because, given the extensive characterization of the *A. thaliana* CLE family over two decades, the likelihood of remaining unannotated canonical CLEs in this specific cluster is minimal. This significantly reduces the risk of ‘label noise’ and provides ideal, high-stringency negative training examples to force the model to distinguish true CLE ‘signal’ from mere structural neighbors.

We used proportionally sized datasets for training and testing: 700 training/314 testing sequences for ESM2, and 550 training/156 testing for ProtT5. ‘Easy negatives’ comprised proteins randomly sampled from distant clusters across all 13 proteomes (n=500 for both models), representing proteins which unambiguously did not belong to the CLE family.

We applied a comprehensive name-based filter using regular expressions to exclude proteins with CLE-related terms from negative sets. The filter matched patterns related to the following gene families: CLV3/ESR (CLE), RGF/CLEL/GLV, CIF1/CIF2, IDA/IDL, CEP, and EPF/EPL. It recognized both the standard abbreviations (like CLE) and the full, written-out names of these families (like CLAVATA).

The held-out test set for both models contained 18 *R. sativus* CLE peptides (positive examples), in addition to hard negatives from *A. thaliana* Cluster 0 not used during training. XGBoost hyperparameters were identical for both models: max_depth=6, learning_rate=0.1, n_estimators=200, subsample=0.8, colsample_bytree=0.8. We used CUDA acceleration. Class imbalance was addressed using the scale_pos_weight parameter. A 20% stratified validation split was used to determine the optimal number of boosting rounds, after which the final model was retrained on the complete training set.

### Novel CLE candidate identification

Each trained classifier was applied to all proteins within Cluster 0. Candidates were stratified by prediction confidence: high confidence (probability ≥0.90), medium confidence (0.70–0.90), and above threshold (≥0.50). To maximize reliability, consensus candidates were defined as proteins achieving high-confidence scores in both ESM2 and ProtT5 pipelines independently. This dual-model approach was chosen to provide orthogonal validation, as the two embedding methods capture different aspects of protein sequence information.

### MEME and FIMO analysis

Motif discovery was performed using the Multiple EM for Motif Elicitation (MEME) suite (Bailey *et al*., 2015). MEME identifies statistically overrepresented sequence patterns by iteratively refining position weight matrices through an expectation–maximization procedure. Motif length was constrained to range between 6 and 50 amino acids, and a maximum of 30 motifs were considered. For the CLE-positive domain, the search for one motif was performed with a constrained length of 12 amino acids. Default parameters were retained for all other settings.

FIMO analysis (Grant *et al*., 2011) was used to search for motifs across ESM2 and ProtT5 datasets (probability ≥0.50). Query motif occurrence was assessed using a p-value threshold of 1×10^−4^. For CLE discovery, FIMO results were classified into p-value ranges corresponding to high (p-value<1×10^−11^), moderate (1×10^−11^≤p-value≤1×10^−9^) and low (p-value>1×10^−9^) sequence similarity.

### Sequence alignment and phylogenetic analysis of peptide motifs

A multiple sequence alignment of the RIPP motifs was generated using CLUSTALW (Thompson *et al*., 1994) and queried against the UniProt database using HMMsearch from the HMMER suite (Potter *et al*., 2018), and filtered with an E-value threshold of 10^−6^.

Their phylogenetic distribution was analyzed by constructing phylogenetic trees using the iPhylo suite (https://iphylo.net/tree/) (Li *et al*., 2025). Phylogenetic tree visualization was carried out using the Phylo.io viewer (https://beta.phylo.io/viewer/) (Robinson *et al*., 2016). Multiple sequence alignment of *P. persica* CLE peptides was performed using Clustal Omega (Madeira *et al*., 2024). Sequence alignment was visualized using Jalview (Waterhouse *et al*., 2009).

### Synthetic CLE peptides

PEP1 (RIT[Hyp]NGANPLHN), PEP2 (RKVYTG[Hyp]NPLHN) and AtCLE27 (AT3G25905.1; RIV[Hyp]SC[Hyp]DPLHN) peptides, with a purity >95%, were commercially synthesized (Bio-Fab Research s.r.l.). Peptides were dissolved to a stock concentration of 1 mM using DMF as solvent and distilled water as diluent. Working solutions were freshly prepared from stock solutions at final concentrations of 1 µmol L^−1^ and 100 nmol L^−1^.

### Tilted-plate assay

*Nicotiana tabacum* seeds were surface-sterilized and sown on half-strength Murashige and Skoog (½MS) medium (Duchefa Biochemie B.V.) supplemented with 1.5% (w/v) sucrose and 1% (w/v) agarose, adjusted to pH 5.7, in square Petri dishes. Synthetic peptides were included in the medium at final concentrations of 1 µmol L^−1^ or 100 nmol L^−1^. Control plates were prepared using the same medium without peptide supplementation. For each peptide treatment, three plates were prepared at the 1 µmol L^−1^ concentration and two plates at the 100 nmol L^−1^ concentration, with eight seeds sown per plate. In addition, five plates without any treatment (no peptide) were included as negative controls. Plates were tilted at a 45° angle with respect to the gravity vector and incubated in a growth chamber at 22 °C under a 16 h light/8 h dark photoperiod for 12 days. Images were acquired after 12 days of growth and morphological parameters were quantified using the SmartRoot tool (Lobet *et al*., 2011) within the ImageJ software (Schneider *et al*., 2012). Statistical analyses were conducted on primary root length data using Tukey’s honestly significant difference (HSD) test (p < 0.05).

## Results

To systematically identify CLE peptides across plant proteomes, we developed a dual-model framework combining pLMs with supervised learning, as outlined in ***Fig. 1A***. Despite their architectural differences, both ESM2 and ProtT5 embeddings revealed remarkably consistent clustering of CLE peptides (***Fig. 2A, B***). ESM2-based Leiden clustering (k=100) identified 165 clusters, while ProtT5-based clustering yielded 152 clusters. In both cases, a single cluster (designated Cluster 0) emerged as strongly enriched for CLE-related sequences (***Fig. 2C, D***), containing 96.7% (ESM2) and 97.3% (ProtT5) of training positives.

**Fig. 2.**
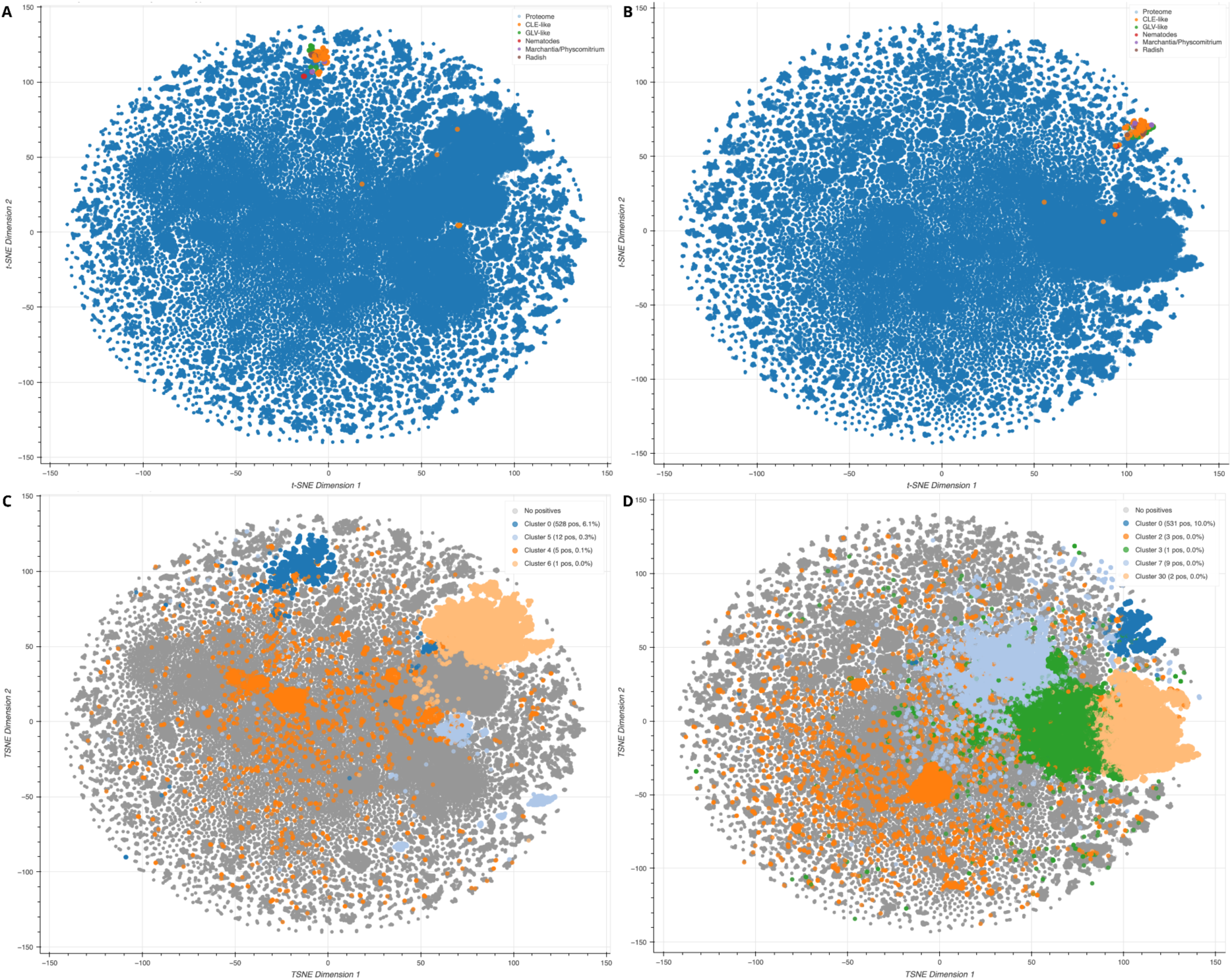
Protein Language Models map CLE peptides into distinct, conserved regions of the proteomic space. **(A-B)** t-SNE visualization of the protein embedding space generated by ESM2 **(A)** and ProtT5 **(B)**. The background proteome is depicted in blue. Colored points represent known signaling peptides: CLE-like (orange), GLV-like (green), nematode CLE mimics (red), basal land plant CLEs (*Marchantia/Physcomitrium*, purple), and radish CLEs (brown). Note that CLE peptides from diverse species cluster tightly together, indicating captured evolutionary constraints. **(C-D)** Identification of CLE-enriched clusters. The plots highlight the specific Leiden clusters containing the highest density of positive training labels against a non-enriched background (gray). In both ESM2 **(C)** and ProtT5 **(D)** spaces, a single primary cluster (“Cluster 0”, blue) captures the vast majority of known CLEs (>96%), effectively isolating the CLE family from the rest of the proteome.

Notably, despite being unsupervised and blind to sequence labels, the clustering naturally grouped the *R.sativus* peptides with known CLEs from other species. This association confirms that both embedding methods capture conserved structural and functional features of the CLE family across different plant lineages. This validates the biological relevance of the feature space, demonstrating its capability to identify CLE candidates even in newly sequenced species lacking prior annotation.

### Classifier performance

Both XGBoost classifiers achieved near-perfect discrimination, with complementary strengths. The ESM2-based classifier achieved AUC-ROC = 1.0000 and 99.4% accuracy on the test set; the ProtT5-based classifier achieved AUC-ROC = 0.9996 and 98.9% accuracy. Notably, both classifiers yielded no false negatives, correctly identifying all 18 *R. sativus* CLEs. The few false positives (two per each model), predicted with high confidence, corresponded to distinct *A. thaliana* proteins: *A0A1P8AMZ2* and *A0A1P8ARH5* (identified by ESM2), and *Q3E7N2* and *P93308* (identified by ProtT5). Despite their current annotation as transmembrane or uncharacterized mitochondrial protein, these candidates may represent structurally similar small signaling peptides. Indeed, it is often observed that lipophilic signal peptides can be misannotated as transmembrane domains (Krogh *et al*., 2001).

As an additional validation, the trained models were applied to proteins within Cluster 0 including in their textual description a term matching the CLE family (these proteins were excluded from both training and candidate sets, as discussed in the Materials and Methods section). For ESM2, 438 proteins were evaluated: 51.6% achieved high confidence, with a probability ≥0.90. In the case of ProtT5, 463 proteins were evaluated, with 54.4% reaching the same threshold. These moderate detection rates do not represent a limit of the classification model. Rather, they reflect the broad scope of our name filter, which retrieved not only CLE peptides but also members of related families (such as CEP, EPF, IDA, and RGF/GLV/CLEL). While these peptides share structural similarities with CLEs, the models correctly distinguished them by assigning lower probabilities. This selectivity is a desirable property, indicating that the classifiers learned CLE-specific features rather than general small-peptide characteristics.

### Novel CLE candidates and consensus selection

Application of the classifiers to unannotated proteins in Cluster 0 identified 410 high-confidence candidates (≥0.90) using ESM2 and 503 using ProtT5. To assess model agreement, we compared candidate sets at different confidence thresholds (***Table 1***). At the most stringent threshold (probability ≥0.99), 294 candidates were identified by both models, representing 94% of ESM2 (313) and 74% of ProtT5 (398) predictions. This substantial overlap increased at lower thresholds, with 85% consensus among all candidates above a 0.5 probability. Candidates were distributed across all 13 proteomes, with the highest numbers in *G. max*, *Z. mays*, *P. vulgaris*, and *L. japonicus*. The majority of proteins were annotated as “Uncharacterized protein” in UniProt. While canonical CLEs are typically <150aa, the classifier identified valid high-confidence candidates up to ∼230aa (Median: ∼100 aa; Range: ∼45–229 aa), potentially representing precursors with longer variable regions or including multiple domains.

**Table 1.** Number of CLE candidates predicted by ESM2- and ProtT5-based classifiers at different confidence thresholds and their overlap. The table reports total candidates identified by each model and the size of their intersection, highlighting increasing consensus at lower probability cutoffs.

| Confidence threshold | ESM2 samples | ProtT5 samples | Intersection |
| --- | --- | --- | --- |
| Top 100 | 100 | 100 | 56 |
| $\geq 0.99$ | 313 | 398 | 294 |
| $\geq 0.90$ | 410 | 503 | 365 |
| $\geq 0.5$ | 540 | 600 | 461 |

The complete list of 365 high-confidence consensus candidates (probability ≥0.90 in both models) is provided in ***Supplementary Table 1***, including UniProt identifiers, species assignment, prediction scores from both models and full protein sequences. Prediction data for all sequences in cluster 0 are available at https://github.com/sales-lab/uncleash.

### Motif discovery and CLE domain validation by MEME and FIMO analyses

We performed a motif discovery analysis using the MEME suite to assess whether the protein candidates we identified shared conserved sequence features characteristic of CLE peptides. Our searches were conducted separately for ESM2- and ProtT5-derived candidates.

To establish a baseline for CLE domain detection, we performed motif discovery on the complete set of known CLE peptides. This analysis identified, as expected, a highly conserved 12-aa motif (***Fig. 3A***) that closely matches the consensus CLE sequence reported in previous studies (Han *et al*., 2016). The resulting position weight matrix was extracted and used as a reference CLE motif for subsequent FIMO analyses.

**Fig. 3.**
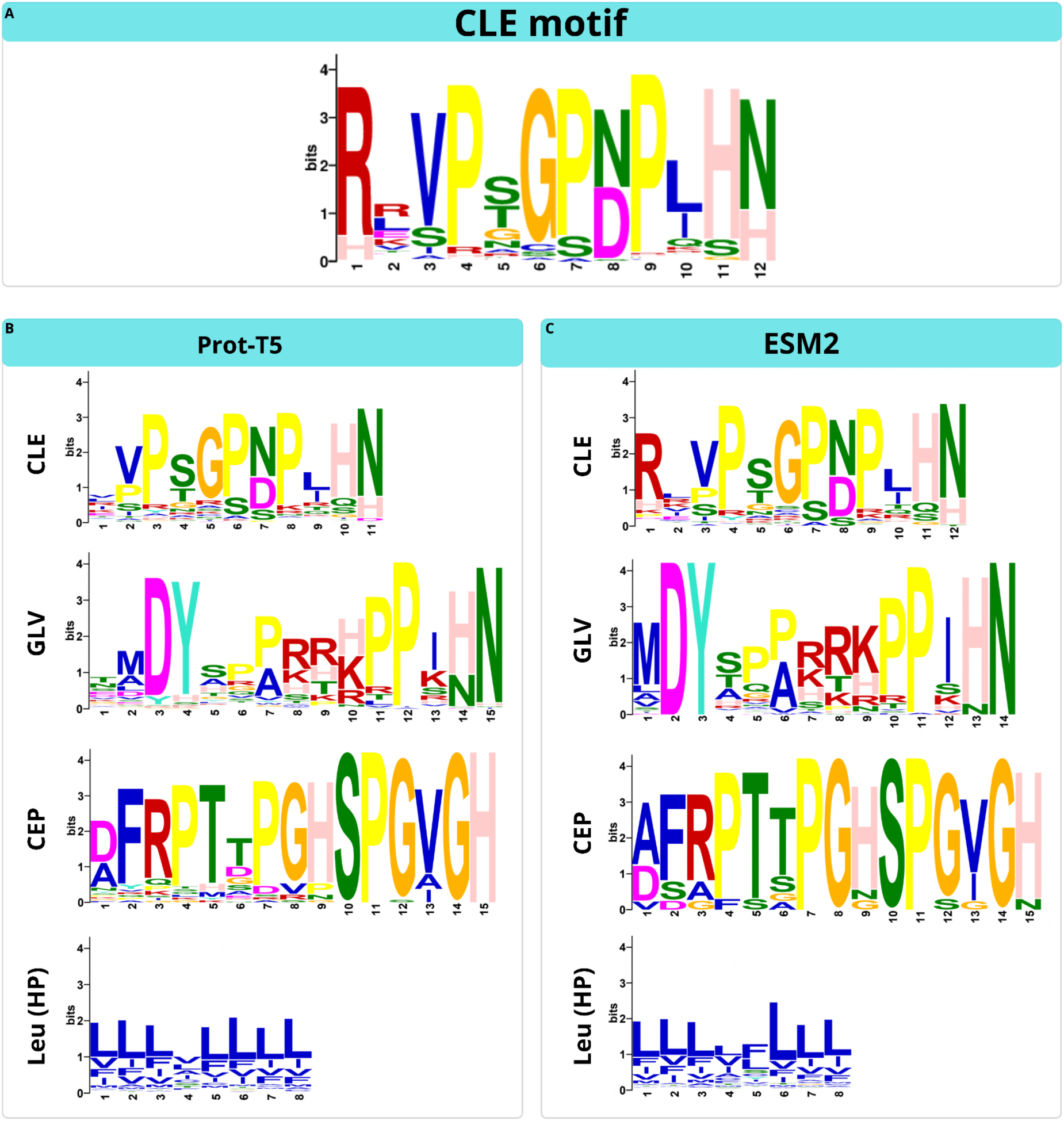
CLE motif and MEME-derived motifs from ESM2 and ProtT5. **(A)** Reference CLE motif generated by MEME using the positive training set of known CLE peptides. This motif was used as the input for FIMO scanning. **(B-C)** *De novo* motif discovery performed on high-confidence candidates identified by ProtT5 **(B)** and ESM2 **(C)**. In both candidate sets, MEME consistently identified the canonical CLE domain (top), as well as motifs characteristic of other signaling families (GLV and CEP). The analysis also captured a conserved hydrophobic leucine-rich motif (Leu HP) typically associated with the N-terminal signal peptide required for secretion.

We then used FIMO to search for CLE motif instances across the ESM2 and ProtT5 datasets. The complete results are reported in ***Supplementary Table 2*** and were subdivided by defining distinct p-value categories. Sequences exhibiting a p-value ≤ 1×10^−11^ were classified as *bona fide* CLE peptides. Based on this stringent threshold, 132 ESM2 candidates (24.4%) and 134 ProtT5 candidates (22.3%) contained a CLE-like motif displaying high sequence similarity to the canonical CLE domain. In addition, 67 ESM2 (12.4%) and 64 ProtT5 (10.7%) sequences showed moderate similarity to the CLE motif, with p-values ranging between 1×10^−11^ and 1×10^−9^. The remaining candidates, which exhibited p-values above this threshold, were classified as having low similarity to the CLE domain (***Supplementary Table 2***).

In parallel, we performed *de novo* motif discovery using MEME. Across all MEME runs, a motif corresponding unambiguously to the canonical CLE domain (∼12 amino acids) was consistently identified as the most significant motif in both ESM2- and ProtT5-derived candidate sets (***Fig. 3B, C***), present in 64.1% of ESM2-high confidence and 59,6% of ProtT5-high confidence (***Supplementary Table 3***).

In addition, MEME identified several recurrent secondary motifs that were shared between the two candidate sets. These included hydrophobic, leucine-rich regions, typically located in the N-terminal portion of the precursor sequences (***Fig. 3B, C***, 100% for ESM2-high confidence and 99% for ProtT5-high confidence), consistent with signal peptide features required for secretion. Notably, motif composition analysis revealed that the AI-driven discovery was not exclusively restricted to CLE peptides. The candidate sets additionally included peptides belonging to related signaling families, most prominently GLV peptides (***Fig. 3B, C***, 14.4% of ESM2-high confidence and 17.5% of ProtT5-high confidence), and CEP (***Fig. 3B, C***, 2.9% of ESM2-high confidence and 8.5% of ProtT5-high confidence), as well as additional small secreted peptides characterized by conserved C-terminal motifs (***Supplementary Table 3***).

The analysis of the sequences harbouring a CLE-motif allowed the identification of a recurring Arg-Ile-Pro-Pro motif in a group of candidate peptides we call RIPP putative peptides. This motif was consistently found with high significance in our sequence analysis (FIMO, MEME). We used these sequences to create a consensus motif, then scanned our datasets again to identify a comprehensive set of RIPP sequences. This resulted in 19 sequences associated to a stringent p-value ≤ 1×10^−15^ (***Supplementary Table 4***), which were further examined through their UniProt entries and found to lack functional annotation. The alignment of these 19 sequences shows classical secreted signal peptide features, like N-terminal hydrophobic targeting signal, a variable region and a conserved C-terminal motif ***(Supplementary Fig. 1***). Collectively, these results establish the presence of a previously uncharacterized group of peptides sharing the RIPP motif (***Supplementary Fig. 2***), providing the foundation for subsequent functional investigations.

### Conservation and evolution of peptide motifs

To further characterize the composition of the CLE-enriched embedding space, we performed *de novo* motif discovery on the union of all candidates identified by either model (probability ≥0.5; n=679 sequences). Beyond the expected CLE domain (51% of sequences) and N-terminal signal peptide (95%), the analysis revealed the presence of related signaling peptide families, as GLV/RGF peptides (15%) and CEP peptides (12%) and other recurrent motifs. Hierarchical clustering of species based solely on the 30 best-ranking motif distribution (presence/absence) patterns (***Fig. 4***) largely recapitulated known phylogenetic relationships. Notably, the use of Jaccard distance (which emphasizes shared motif presence while ignoring shared absences) resulted in a cleaner separation of deep evolutionary lineages, correctly placing bryophytes as an outgroup and clustering angiosperm families according to their shared motif repertoires. Bryophytes (*M. polymorpha* and *P. patens*) exhibited few motifs in common with other species, indicating their evolutionary independence. Fabaceae (*G. max*, *L. japonicus*, and *P. vulgaris*) formed a tight cluster. Rosaceae (*F. vesca* and *P. persica*) shared some motifs with the Fabaceae consistent with their common membership among Rosids.

**Fig. 4.**
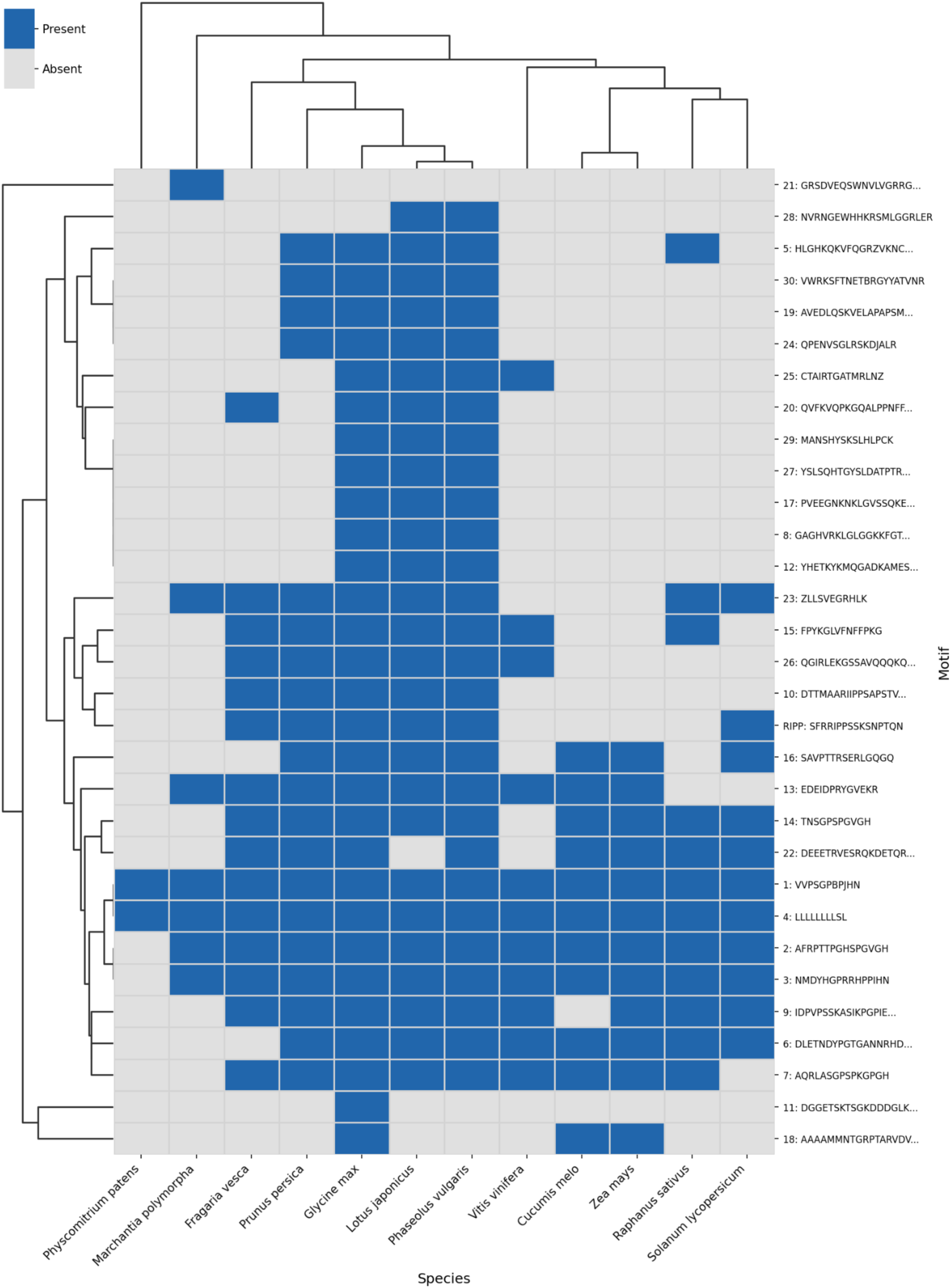
Motif distribution across plant species in the CLE-enriched embedding space. Binary heatmap showing the presence (blue) or absence (gray) of MEME-identified motifs across 12 plant proteomes. Hierarchical clustering was performed using the Average Linkage (UPGMA) method based on Jaccard distances to minimize the weight of shared absences. Clustering was applied to both motifs (rows) and species (columns). The analysis includes candidates from both ESM2 and ProtT5 models (probability ≥0.5). Motifs are labeled by their MEME identifier and consensus sequence. MEME-1 (VVPSGPBPJHN) corresponds to the CLE domain; MEME-4 (LLLLFLLL) represents the N-terminal signal peptide; MEME-3 (TMDYHPPRRHPPIHN) matches the GLV family; MEME-2 (DFRPTTPGHSPGVGH) corresponds to CEP peptides. The RIPP motif is shown separately. In MEME consensus sequences, ambiguous residue codes are used: B represents Asx (Asp or Asn), J represents Xle (Leu or Ile) and Z represents Glx (Glu or Gln).

Single members of other plant families were also separated into distinct clusters, including Solanaceae (*S. lycopersicum*), Vitaceae (*V. vinifera*), Brassicaceae (*R. sativus*), and Poaceae (*Z. mays*). This pattern is largely consistent with evolutionary relationships captured by the embedding space, although quantitative concordance testing is required. The co-occurrence of CLE, GLV, and CEP peptides within the same embedding neighborhood likely reflects their shared biochemistry and their evolutionary relationships, rather than strict sequence homology, reinforcing the concept of ‘distributed peptide-ness’ captured by pLMs.

In order to explore the evolutionary distribution of RIPP putative peptides, a motif-based search against the UniProt database (*version 2025_01*) identified RIPP-related sequences across a broad range of plant species albeit limited to Magnoliopsida (***Supplementary Fig. 3***). Matches were detected with high statistical confidence (E-value < 10^−6^), indicating that RIPP-related sequences are conserved beyond a limited set of taxa. The identified peptides span multiple but not all Magnoliopsida lineages, supporting a widespread evolutionary distribution of RIPP motifs with probable losses, as for Brassicaceae, thus confirming the absence seen in *R. sativus* (***Fig. 4***). The taxonomic distribution across plant families of RIPP putative peptides is summarized in ***Supplementary Fig. 3*.**

### From Prediction to Experiment: Selecting CLE Peptides for In Vitro Studies

To prioritize candidates for experimental validation (***Fig. 1C***), we focused on peptides consistently ranked as top hits by both models. Two high-confidence candidates were selected based on their consensus ranking, presence of the conserved CLE domain, and biological relevance.

The first candidate (PEP1: RITPNGANPLHN) was identified in *P. persica* (A0A251Q718) and represents a previously unreported CLE peptide. Alignment of PEP1 with other known CLE peptides from *P. persica* is provided in ***Supplementary Fig. 4***. Notably, PEP1 was supported by high-confidence scores from both embedding-based models (ESM2: 91°, ProtT5: 121°) and exhibits some similarities to SlCLE38 (Carbonnel *et al*., 2022).

The second candidate (PEP2: RKVYTGPNPLHN) was identified in *P. vulgaris* (V7CV52) and showed exceptional agreement between the two models, ranking within the top tier in both ESM2 and ProtT5 predictions (12° and 15°, respectively). While this and other CLE-like sequences with this motif have been previously identified in *P. vulgaris* (PvCLE20, (Hastwell *et al*., 2015)) and in a limited number of species, this peptide has not been functionally characterized, nor has its biological activity been assessed *in vivo*. Sequence alignment revealed close similarity to SlCLE45 and AtCLE22, suggesting potential conservation of function. Notably, homologous *CLE* genes with identical peptide sequence were also detected in agronomically relevant and experimentally challenging species, such as *Olea europaea* (olive tree), which further motivated its selection for experimental investigation.

Both PEP1 and PEP2 were predicted as CLE peptides by the XGB classifiers with scores exceeding 0.999.

An easy way to test potential peptide activity is to assess the effects of candidates on seedling root development (Yamaguchi *et al*., 2016; Fletcher, 2020; Stührwohldt *et al*., 2020). Tilted-plate assays were carried out in *N. tabacum* seedlings which exhibited distinct and peptide-specific root growth responses (***Fig. 5***). Treatment with PEP2 resulted in a reduction in primary root length compared to untreated controls, clearly observable already at 100 nmol L^−1^ and more pronounced at 1 µmol L^−1^ (p < 0.05; ***Fig. 5***). In contrast, PEP1-treated seedlings displayed root lengths comparable to those of control seedlings, indicating a lack of measurable effect on root elongation under the same experimental conditions; therefore, further analyses will be needed to assess its biological activity. As expected, treatment with AtCLE27 caused a strong inhibition of root growth. Together, these results demonstrate that PEP2, but not PEP1, negatively regulates root elongation in *N. tabacum*, consistent with the bioactivity reported for other CLE peptides.

**Fig. 5.**
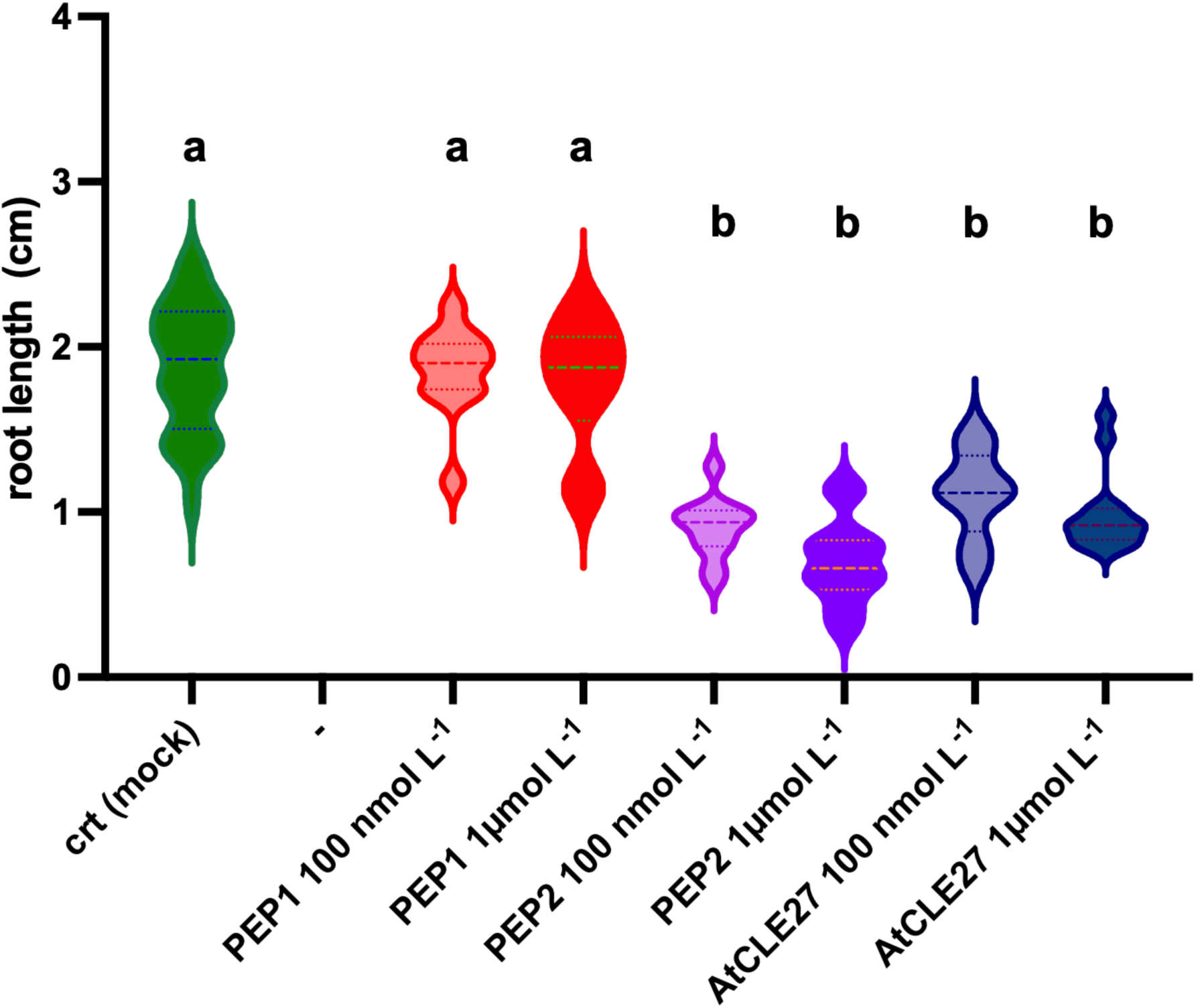
Effect of different CLE peptides on the primary root length of tobacco seedlings. Violin plots show the distribution of total root length (cm) measured after 12 days of growth on vertical half-strength MS (½ MS) plates supplemented with either 100 nmol L^−1^ or 1 µmol L^−1^ of PEP1 and PEP2. Crt indicates untreated control seedlings grown on ½ MS medium without peptide supplementation. The Arabidopsis peptide AtCLE27 was used as a positive control. The width of each violin represents the probability density of the data at different values. Inside each violin, the dashed horizontal line represents the median, and the dotted lines represent the upper and lower quartiles (25th and 75th percentiles). Whiskers extend to the minimum and maximum data points within 1.5 times the interquartile range (IQR). The width of the plots is scaled by the number of observations (n). Groups sharing the same letter are not significantly different (Tukey’s HSD test, p < 0.05). n = 8-30 seedlings per treatment.

## Discussion

### AI-driven embeddings capture conserved CLE features beyond sequence similarity

Artificial intelligence is revolutionizing biological research, and plant science is no exception. Deep learning has revolutionized protein sequence analysis, revealing hidden layers of biological complexity that conventional methods cannot access. Our work exemplifies this transformation. We leveraged pLM trained on evolutionary-scale sequence data to develop a framework capable of identifying previously undetected signaling peptides.

Peptide-mediated signaling is essential for plant development, yet many conserved peptide families remain incompletely characterized due to low primary sequence conservation outside short functional motifs. CLE peptides exemplify this challenge: despite being evolutionarily ancient and functionally critical, a substantial fraction of family members remains uncharacterized due to functional redundancy and the minimal length of the conserved C-terminal domain. Here, we applied an AI-driven framework combining pLM with supervised learning to systematically identify CLE peptides across 13 plant proteomes, revealing previously unannotated candidates.

A central finding of this study is the remarkable concordance between two architecturally distinct pLMs. ESM2 and ProtT5 were trained with different objectives, process sequence information through fundamentally different computational strategies, and generate embeddings of different dimensionality. Yet both models independently converged on essentially the same CLE-enriched cluster. The substantial overlap observed across confidence thresholds further demonstrates that these models are capturing the same underlying biological signal.

This convergence is not coincidental but reflects the existence of evolutionarily constrained sequence features within CLE precursors (Gentile *et al*., 2025) that both models have learned to recognize. Because these models were trained on hundreds of millions of protein sequences spanning all domains of life, they have internalized statistical regularities imposed by millions of years of evolution. When two such models, operating through fundamentally different computational strategies (Elnaggar *et al*., 2022; Lin *et al*., 2023), independently identify the same proteins as CLE-related, this constitutes strong evidence that these sequences share genuine biological properties rather than superficial sequence artifacts. The cross-species nature of our analysis further supports this interpretation. As evident in ***Fig. 2A*** and ***B***, CLE peptides from various plant lineages cluster together in embedding space, indicating functional constraints that have persisted since the last common ancestor of land plants. Notably, GLV peptides and parasitic nematode CLE mimics also cluster in close proximity to the plant CLEs. This convergence, spanning deep evolutionary time and distinct biological kingdoms, underscores the fact that pLM embeddings accurately capture the biological essence of CLE identity, rather than merely reflecting statistical patterns specific to particular lineages.

Traditional approaches to CLE identification have focused almost exclusively on the canonical 12-residue C-terminal motif. While this motif constitutes the functional core of mature peptides, our results reveal that the CLE identity extends far beyond this short conserved sequence. By analyzing entire pre-propeptides rather than isolated motifs, pLM captures what we term “distributed CLE-ness“: a constellation of sequence features that collectively define family membership.

These distributed features likely encompass multiple functional elements: (i) N-terminal signal peptides with characteristic hydrophobic profiles required for secretion into the apoplast; (ii) variable regions with compositional biases influencing processing efficiency or precursor stability; (iii) sequence contexts flanking the CLE domain that modulate proteolytic cleavage specificity; (iv) subtle sequence signatures inherited from ancestral genes through the duplication events that expanded this family; (v) and finally the CLE domain. Crucially, many of these features may be undetectable through conventional alignment because they represent statistical tendencies rather than strictly conserved positions. This perspective fundamentally reframes CLE peptide identification. Rather than simply asking “does this sequence contain a recognizable CLE motif?”, the embedding-based approach implicitly asks “does this sequence occupy the same region of protein space as known CLE precursors?”. In the learned representation, functionally related proteins cluster together regardless of overall sequence identity, while functionally divergent proteins occupy distant regions. This shift from motif-centric to holistic sequence analysis enables detection of divergent family members that retain functional CLE properties despite lacking recognizable similarity to canonical members.

A distinctive feature of our approach is the systematic focus on proteomes rather than genomes. Previous large-scale CLE surveys have primarily mined genomic sequences, searching for open reading frames encoding the characteristic CLE domain (Han *et al*., 2016; Goad *et al*., 2017; Carbonnel *et al*., 2022). While valuable, genome-based approaches face challenges including gene prediction errors, alternative splicing complexity, and the gap between genomic potential and actual protein expression. By operating directly on annotated proteome sequences, our analysis captures proteins that are actually expressed and recognized by annotation pipelines, providing a more direct connection to biological function. The proteome-wide perspective also enables systematic comparison across species at unprecedented scale. By embedding proteins from diverse plant species into a common feature space, we can directly assess how CLE-related sequences from different lineages relate to one another, revealing patterns of deep conservation that persist across broad evolutionary distances.

An important methodological consideration concerns the candidate filtering step, which relies on UniProt annotations. As a consequence, peptides lacking formal UniProt annotation may appear among the predicted candidates even if they have been previously proposed in the literature as CLE peptides. A clear example is provided by several tomato CLE peptides reported by Carbonnel *et al*. (2022), some of which are recovered in our output list because they were not being annotated as CLE in public databases. In this sense, our results build upon prior work identifying these candidates, and their recovery reflects annotation gaps rather than biological novelty. Accordingly, when applying this framework to the analysis of individual proteomes, careful comparison with the literature, assessment of the most recent annotations, and appropriate tuning of filtering criteria will be essential to achieve the desired level of biological resolution.

The MEME suite analyses support this interpretation. Crucially, motif analysis was applied only downstream of candidate selection, serving as an interpretative tool rather than a discovery constraint. FIMO analysis allowed us to scan the sequences and confirmed the presence of a notable number of currently unannotated CLE sequences, with 132 ESM2 candidates (24.4%) and 134 ProtT5 candidates (22.3%) showing excellent p-value scores. Moreover, *de novo* motif discovery using MEME enabled us to dissect the composition of the predicted peptide space. In this context, the CLE motif emerges not as the primary driver of discovery, but as a final discriminator that helps classify and rationalize the biological identity of the recovered sequences. As expected, CLE domain emerged as the dominant motif across all candidate sets; alongside this, secondary motifs corresponding to signal peptides and other conserved regions were also consistently identified. Moreover, the candidate sets included not only CLE peptides but also members of related signaling families such as GLV, CEP and related small secreted peptides. This indicates that the embedding framework captures shared biochemical and structural properties of the broader secreted peptide signaling families rather than relying solely on the CLE core motif.

Concerning the non-peptide MEME-motifs, the identification of conserved sequence stretches across candidate peptides provides an additional layer of interpretation to embedding-based analyses. Rather than representing isolated biochemical features, these motifs likely act as evolutionary signatures that have been retained under selective pressure. In this framework, motif composition emerges as a proxy for the evolutionary history of peptides, capturing deep conservation features. The concordance between motif-based signatures and embedding-derived relationships supports the view that pLM integrate evolutionary constraints, reinforcing the notion that conserved motifs, beyond the canonical peptide motif, constitute distributed evolutionary fingerprints within peptide families, offering insight into how these signaling elements have evolved.

Interestingly, our analyses identified a subset of sequences belonging to a broader class of peptides that we termed as RIPP putative peptides. These sequences were detected in the FIMO analysis with low p-value scores and were misclassified by MEME as CLE motifs. This observation places them within a wider sphere of small signaling peptides that are not *bona fide* CLEs but are frequently recognized (and in some cases misannotated) as such. These candidates lacked UniProt functional annotation, suggesting that they remain largely uncharacterized. HMMsearch analysis revealed that putative RIPP peptides are present across a wide range of Magnoliopsida. Nevertheless, they were not detected in specific families, as in the Brassicaceae. Notably, in the majority of cases, these sequences are either uncharacterized or misannotated as transmembrane proteins, and only in rare and sporadic instances as CLE-like peptides. Nevertheless, although these peptides display partial similarity to CLE family members, they cannot be classified as canonical CLE peptides due to their atypical motif. In particular, a primary indicator is the mispositioning of proline residues, which are critical determinants of CLE peptide bioactivity and must occupy highly conserved positions (Jun *et al*., 2007). This deviation strongly supports a distinct functional identity. We therefore speculate that RIPP sequences may represent a previously uncharacterized class of plant signaling peptides. The identification of RIPP putative peptides highlights the model’s ability to group peptides based on structural mimicry. While distinct from CLEs, their consistent prediction of a conserved structural topology typical of a secreted peptide (i.e. a signal sequence with a hydrophobic portion and a conserved motive at the C-terminal flanked by a cleavage sites) suggests they are *bona fide* secreted signaling peptides, possibly representing a novel target for future functional studies.

Any computational methodology ultimately requires experimental validation to demonstrate its biological relevance. In this context, computational approaches are particularly powerful in increasing experimental efficiency, enabling more targeted hypotheses and reducing the need for large-scale, untargeted screening. Here, we presented a fast and scalable computational framework and complemented it with an *in vivo* application, illustrating how the prioritization of a limited number of candidates can nevertheless lead to biologically meaningful outcomes. In particular, we selected a completely novel CLE peptide (PEP1) and a previously reported but functionally unexplored CLE-like sequence (PEP2) that enabled us to combine sequence novelty with biological relevance, providing complementary insights into CLE peptide diversity and function across plant species. By testing these two peptides, we showed that PEP2 from *P. vulgaris* significantly reduced primary root growth, which is widely used as a proxy for CLEs bioactivity (Yamaguchi *et al*., 2016; Fletcher, 2020; Stührwohldt *et al*., 2020). To our knowledge, PEP2 had not been functionally characterized before, making this observation both novel and coherent with previously explored CLEs activities. Interestingly, PEP2 differs from the closely related tomato peptide SlCLE45 by a single amino acid R-to-Y substitution at position 4, usually occupied by a proline residue (***Fig. 3A***). Notably, SlCLE45 was reported to have no effect on root growth in tomato (Carbonnel *et al*., 2022), suggesting that subtle sequence variation at this position may have a substantial impact on biological activity, regardless of the absence of the conserved P. In contrast, PEP1 did not exhibit measurable effects on root growth under the conditions tested. Further analyses will be required to elucidate the biological role of this peptide, including roles that may manifest outside the experimental conditions tested here. Overall, these findings underscore the value of combining *in silico* predictions at evolutionary scale with targeted *in vivo* assays on a model species, and highlight how such integrative strategies can accelerate the discovery of functional signaling peptides.

### Scalability of the framework, limits, future perspectives

The framework presented here is inherently scalable. Thanks to the efficiency of modern pLMs and the use of high-performance computing, the embedding and classification of entire proteomes can be completed in a matter of hours. This stands in stark contrast to the weeks or months required for the experimental characterization of even a single peptide candidate.

Consequently, the primary bottleneck in peptide discovery shifts from identification to validation. While our AI pipeline drastically narrows down the search space by prioritizing high-confidence candidates (probability ≥0.90), it does not replace the need for biological verification but rather acts as a strategic guide to focus resources.

A key strength of the presented framework is its inherent generalizability. Although we focused on CLE peptides as a well-defined and biologically relevant case study, the methodology is not restricted to this family. The pipeline requires only two inputs (a set of proteomes to screen and a curated collection of positive examples), and it can be readily extended beyond plant signaling peptides to any protein family for which a reliable positive training set is available. Despite these strengths, several limitations should be acknowledged. The quality of predictions depends critically on the composition and curation of the positive training set. Families with few characterized members, high internal diversity, or poorly defined boundaries may yield less reliable classifiers. In our case, the availability of hundreds of CLE sequences across multiple species provided a solid foundation; however, extending the approach to less-characterized peptide families may require careful dataset assembly and iterative refinement.

In this context, our study demonstrates that protein language models can effectively recover CLE peptides across diverse plant proteomes, overcoming the intrinsic limitations of homology-based methods and genome annotation errors. This computational framework offers unprecedented speed and scalability, allowing the screening of entire proteomes in a fraction of the time required by traditional methods, provided adequate computational resources are available. Importantly, our analysis revealed that the CLE-enriched embedding space encompasses not only canonical CLE peptides but also related signaling families, including GLV and CEP peptides, as well as potentially novel classes such as the RIPP putative peptides described here. This finding underscores the capacity of pLM-based approaches to capture the broader landscape of small secreted peptide signaling in plants. The experimental validation of PEP2, a CLE peptide shared, among others, by *P. vulgaris* and *O. europaea*, confirms that AI-driven predictions can successfully guide the identification of biologically active signaling molecules. While the framework was developed using CLEs as a case study, it is readily applicable to other peptide families for which a curated positive training set is available. Experimental validation remains essential: computational predictions, however robust, require functional confirmation to establish biological relevance. By serving as a high-throughput filter to prioritize candidates, our AI-driven approach effectively bridges the gap between *in silico* prediction and *in vivo* function, accelerating the discovery of novel signaling molecules and expanding the known repertoire of plant peptide hormones.

Looking ahead, the convergence of AI-driven sequence analysis with high-throughput experimental platforms holds transformative potential for plant biology. As pLM continues to improve (Du *et al*., 2025, Preprint) and proteome databases expand, the systematic discovery of signaling peptides and other functional protein classes will become increasingly feasible. This work represents an early contribution to this emerging paradigm, demonstrating that AI-based approaches can complement and extend traditional methods, accelerating the pace of discovery in plant signaling research.

## Supplementary data

Supplementary Table S1: Complete list of 365 high-confidence consensus candidates (probability ≥0.90) resulting from both ESM2 and ProtT5 models.

Supplementary Table S2: Comparative FIMO analysis of peptide candidates derived from ESM2 and ProtT5 embeddings.

Supplementary Table S3: Comparative MEME motif analysis of high-confidence peptide candidates derived from ESM2 and ProtT5 embeddings.

Supplementary Table S4: FIMO identification of candidate RIPP peptides.

Supplementary Fig. S1: Multiple sequence alignment of 19 RIPP putative peptides sequences.

Supplementary Fig S2. RIPP motif.

Supplementary Fig. S3: Phylogenetic distribution of RIPP motif across plants.

Supplementary Fig S4. Multiple sequence alignment of PEP1 with known peach CLE peptides.

## Acknowledgements

This work was supported in part by the high-performance computing infrastructure developed under the project “CONVECS”, funded by the PR Veneto FESR 2021-2027 program, Priority 1 – Specific Objective 1.1 – Action 1.1.2

## Author Contribution

MB, MRN, CF, GS and LT: conceptualization; MRN and GS: methodology; MB, MRN, CF, GS and LT: formal analysis; AP: investigation; GS and LT: resources; MRN: data curation; MB, CF and MRN: writing - original draft; MB, MRN, CF, GS and LT: writing - review & editing; MB, MRN, and CF: visualization; GS, and LT: supervision; LT: funding acquisition.

## Data Availability

The Python code developed for this study, including scripts for embedding generation, clustering, and the XGBoost classification pipeline, is openly available at https://github.com/sales-lab/uncleash. All datasets generated and analyzed during this study are deposited in the same repository. This includes the FASTA files containing the known CLE sequences used in this study, as well as the full lists of identified CLE candidates in FASTA format. The raw proteome datasets analyzed in this study are available from the UniProt database (https://www.uniprot.org/), Phytozome (https://phytozome-next.jgi.doe.gov/), and the Genome Database for Rosaceae (GDR; https://www.rosaceae.org/) using the specific accession numbers detailed in the Materials and Methods section.

